# Conserved transcriptional programming across sex and species after peripheral nerve injury predicts treatments for neuropathic pain

**DOI:** 10.1101/2022.05.30.494054

**Authors:** Shahrzad Ghazisaeidi, Milind M. Muley, YuShan Tu, Mahshad Kolahdouzan, Ameet S. Sengar, Arun K. Ramani, Michael Brudno, Michael W. Salter

**Affiliations:** Department of Physiology, University of Toronto; Toronto, Canada; Program in Neuroscience & Mental Health, Hospital for Sick Children; Toronto, Canada; Centre for Computational Medicine, Hospital for Sick Children; Toronto, Canada; Department of Computer Science, University of Toronto; Toronto, Canada

**Keywords:** Peripheral Nerve Injury, Transcriptomic, Spinal cord, therapy

## Abstract

Chronic pain is a devastating problem affecting 1 in 5 individuals around the globe, with neuropathic pain the most debilitating and poorly treated type of chronic pain. Advances in transcriptomics and data mining have contributed to cataloging diverse cellular pathways and transcriptomic alterations in response to peripheral nerve injury but have focused on phenomenology and classifying transcriptomic responses. Here, with the goal of identifying new types of pain-relieving agents, we compared transcriptional reprogramming changes in the dorsal spinal cord after peripheral nerve injury cross-sex and cross-species and imputed commonalities, as well as differences in cellular pathways and gene regulation. We identified 93 transcripts in the dorsal horn that were increased by peripheral nerve injury in male and female mice and rats. Following gene ontology and transcription factor analyses, we constructed a pain interactome for the proteins encoded by the differentially expressed genes, discovering new, conserved signaling nodes. We interrogated the interactome with the Drug-Gene database to predict FDA-approved medications that may modulate key nodes within the network. The top hit from the analysis was fostamatinib, the molecular target of which is the non-receptor tyrosine kinase Syk, which our analysis had identified as a key node in the interactome. We found that intrathecally administrating the active metabolite of fostamatinib, R406, significantly reversed pain hypersensitivity in both sexes. Thus, we have identified and shown the efficacy of an agent that could not have been previously predicted to have analgesic properties.

**One sentence summary:** Unbiased approach to predicting safe therapies for neuropathic pain

## INTRODUCTION

Chronic pain affects 16-22% of the population and is one of the major silent health crises affecting physical and mental health (*1, 2*). Neuropathic pain, which results from damage to the somatosensory system in the peripheral or in the central nervous system (CNS) (*3*), is the most recalcitrant type of chronic pain. Therapeutic options for neuropathic pain are limited by poor efficacy, side effects, and tolerability of even approved pain medications (*4, 5*).

Damage to peripheral nerves is known to produce persistent functional reorganization of the somatosensory system in the CNS (*6*). The primary afferent neurons in peripheral nerves project into the dorsal horn of the spinal cord, which is the critical first site in the CNS for integrating, processing, and transmitting pain information. Transcriptomic changes in the dorsal horn produced by peripheral nerve injury have been increasingly described (*7–10*) with a large emphasis on characterizing sex differences in changes in gene expression. Such studies are touted to hold promise to characterize pathological biochemical pathways that might in the future reveal targets for new therapies. However, there has been a focus on cataloging transcriptomic changes, unconnected from identifying pain therapeutics. Thus, there remains a gap between describing molecular changes in the dorsal horn and identifying new therapeutics.

Here, we took on the challenge of filling this gap by using a purposeful approach to explore the possibility of identifying pain-relieving drugs in an unbiased way through connecting transcriptomic changes to drug discovery. To gain power in our study we simultaneously looked not just between sexes in a single species but between sexes in two species. Unexpectedly, given the growing prominence of sex differences across biomedical sciences, we found many more commonalities than differences between sexes and across species in the gene expression changes produced in the spinal dorsal horn ipsilateral to the peripheral nerve injury. From the commonalities, we built a species-conserved sex-conserved pain interactome network. With an unsupervised approach, we used this interactome to predict safe therapies that may have the most impact.

## RESULTS

We evaluated dorsal horn transcriptomes after spared nerve injury (SNI), a widely used model of peripheral neuropathic pain (*11*), or sham surgery in male and female mice and rats, seven days surgery (Fig. 1A). We collected gray matter from the dorsal horn of L4-L5 spinal cord ipsilateral and contralateral to the surgery. In order to obtain sufficient RNA from mice, each sample was pooled from two animals. The experimenters who did the dissections, tissue removal and extraction of the RNA were unaware of which animals had undergone SNI or sham surgery.

**Fig. 1.**
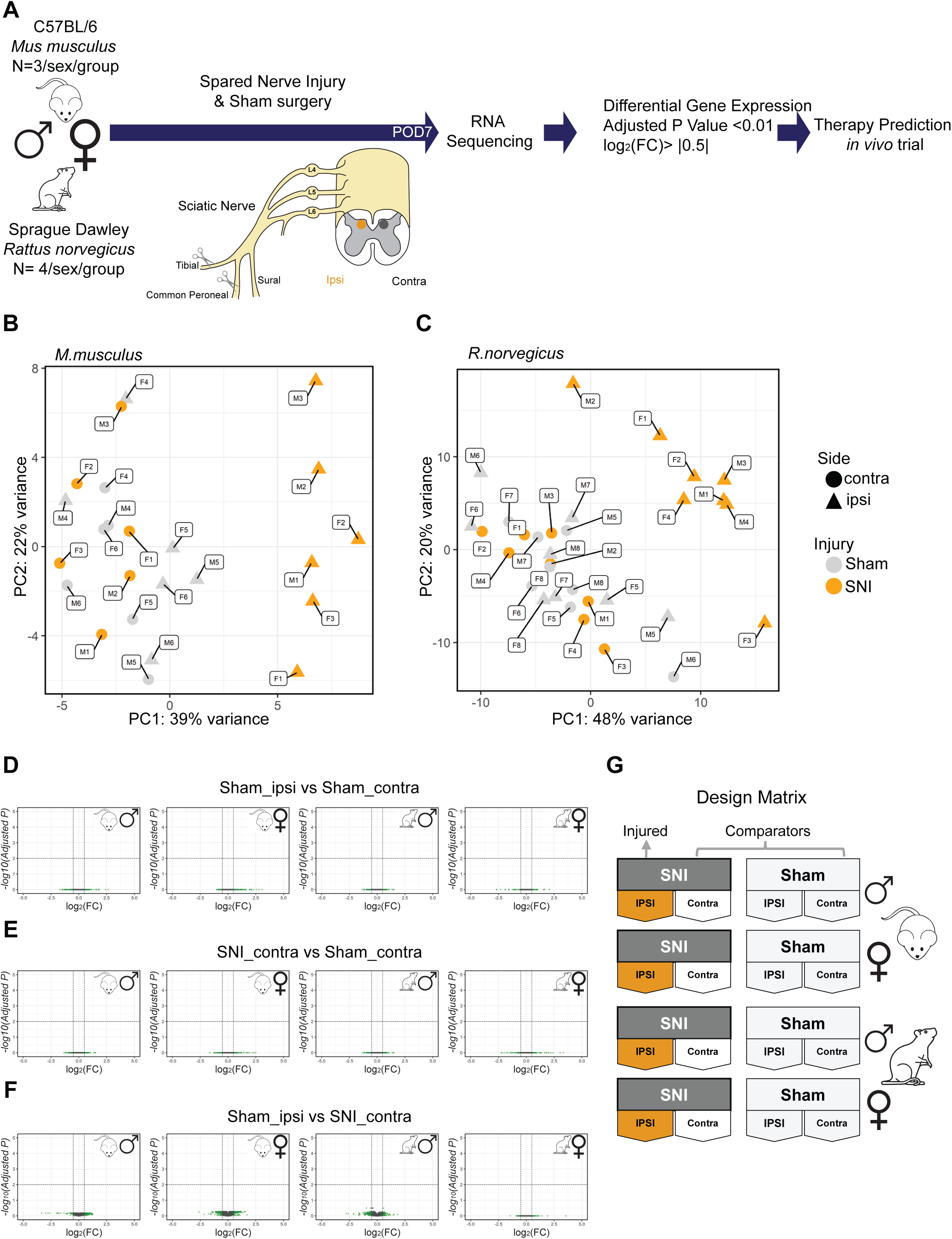
Experimental Design and data overview. (A) The experimental workflow is illustrated. (B-C) Scatter plot representing principal component analyses of the dimensions PC1 versus PC2 samples. (B) Principal component analysis in mice. (C) principal component analysis in rats. (D-F) Volcano plots showing pair-wise differential gene expression in male and female mice and rat between comparators. (D) Sham_ipsi vs Sham_contra. (E) SNI_contra vs Sham_contra. (F) Sham_ipsi vs SNI_contra. (G) Summary of all groups in design matrix.

### Dorsal horn transcriptomes ipsilateral to SNI form a distinct cluster

To define main sources of transcriptome variability we first analyzed the datasets at the sample level by principal component analysis separately for mice and for rats. In the mouse dataset, the two principal components (PCs) PC1 (39%) and PC2 (22%) were the major PCs, explaining 61% of the overall variance (fig. S1). We found in both males and females that there was a clear clustering of the samples from the ipsilateral dorsal horn of animals that had received SNI (SNI_ipsi) as compared to the remaining groups – contralateral to SNI (SNI_contra), ipsilateral to sham surgery (Sham_ipsi), and contralateral to the sham (Sham_contra) (Fig. 1B). In the rat dataset, two principal components explained 48% and 20% of the variance and SNI_ipsi samples were distinct from the remainder (Fig. 1C). Thus, in both species there is a clear cluster primarily across PC1 of SNI_ipsi samples separate from Sham_contra, Sham_ipsi and SNI_contra.

In order to identify differentially expressed genes (DEGs) we did pairwise comparisons between samples of the levels of individual genes. DEGs were defined with the criteria of the adjusted P-value <0.01 and log_2_ fold-change absolute value greater than 0.5 (|log FC|> 0.5). In comparing SNI_ipsi to the other groups we found numerous DEGs (fig. S2) whereas no DEGs were detected comparing Sham_ipsi versus Sham_contra within sex and species (Fig. 1D). Furthermore, there were no DEGs comparing SNI_contra compared with Sham_contra (Fig. 1E) or with Sham_ipsi (Fig. 1F). Taking these findings together we conclude from both principal component analysis and DEG analyses that the transcriptomes of the dorsal horn ipsilateral to SNI are distinct from those contralateral to SNI or either of the sham groups. Moreover, because we found no differences at the gene expression level in SNI_contra, Sham_ipsi and Sham_contra, we combined these groups, by sex or by species, as comparators for the remainder of our analyses (Fig. 1G).

### High correlation in the gene expression pattern of spinal cord dorsal horn in male and female mice and rats after peripheral nerve injury

From this analysis, we compared gene expression levels by sex and by species in the ipsilateral dorsal horn after SNI with those in the respective comparator group (Fig. 2A-D). In mice we observed that after SNI there was increased expression of 278 and 136 genes in males and females, respectively (Fig. 2A, B). In females we found 14 genes expression of which was significantly decreased after SNI where none was decreased in males (Fig. 2A, B & fig. S3). In rats, 271 genes were upregulated in the SNI_ipsi males versus the comparator group and none was decreased. Whereas in females, the expression level of 403 genes was increased, and that of 13 genes was decreased, after SNI (Fig. 2C-D & fig. S3). Thus, in both mice and rats downregulation of gene levels after SNI was only observed in females, and of these DEGs there were 4 in common in both species.

**Fig. 2.**
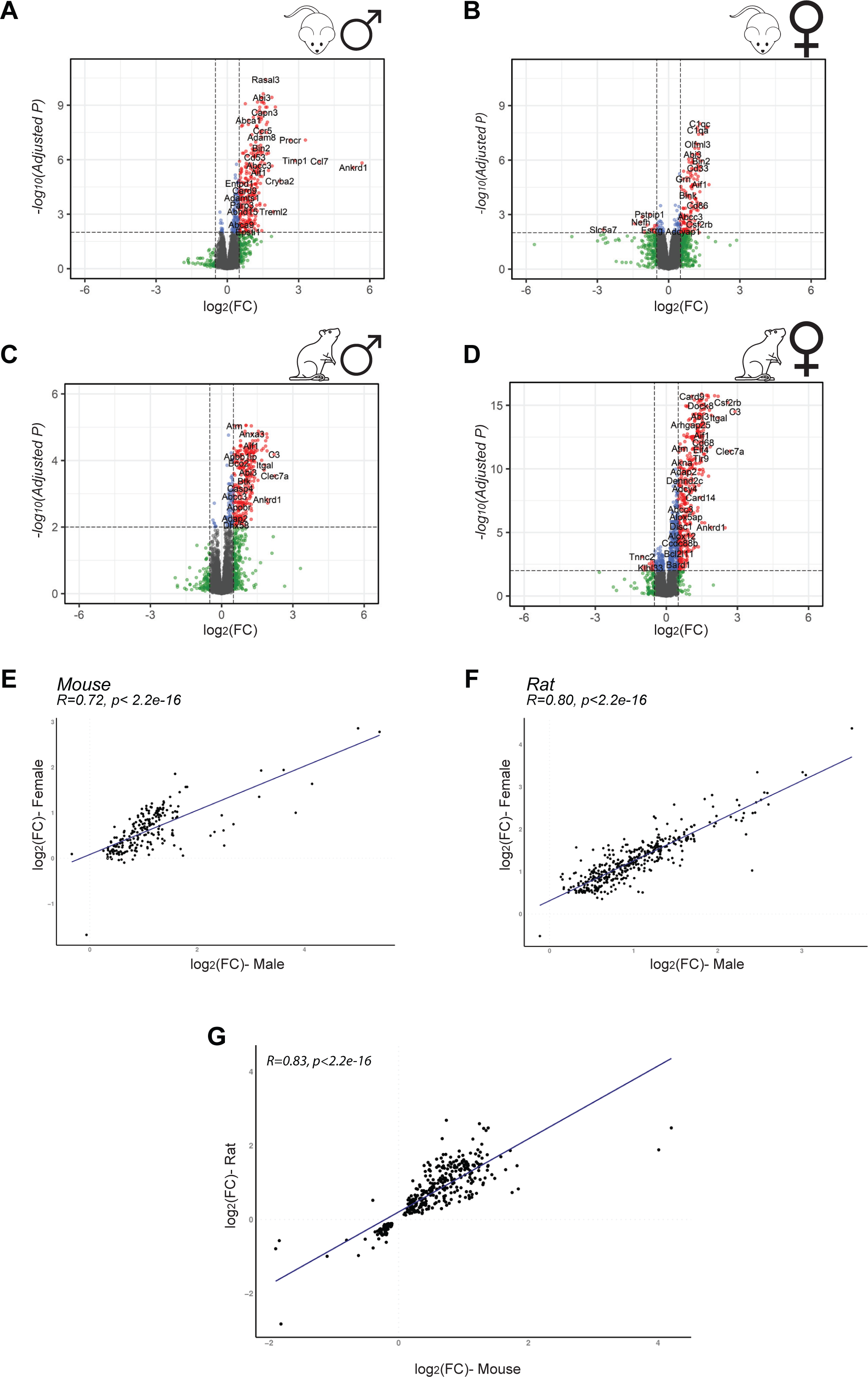
Transcriptome changes after SNI in male and female rodents. (A-D) Volcano plots were obtained by plotting the log2 fold change of SNI_ipsi against the negative Log10 of the EdgeR adjusted p-value. Genes that changed 0.5 log2(FC) or more with a significance of adj p-value <0.01 are shown red. Genes that were differentially expressed significantly (p < 0.01) but changed less than 0.5 log2(FC) are highlighted in blue and black dots are insignificant changes. (A) male mice, (B) female mice, (C) male rat and (D) female rat. (E-G) Linear correlation of log2(FC) of SNI_ipsi vs comparators is demonstrated. Genes that were differentially expressed in at least one dataset is considered. (E) Pearson correlation in male and female mice. (F) Pearson correlation between male and female rat. (G) and the Pearson correlation between mice and rat.

**Fig. 3.**
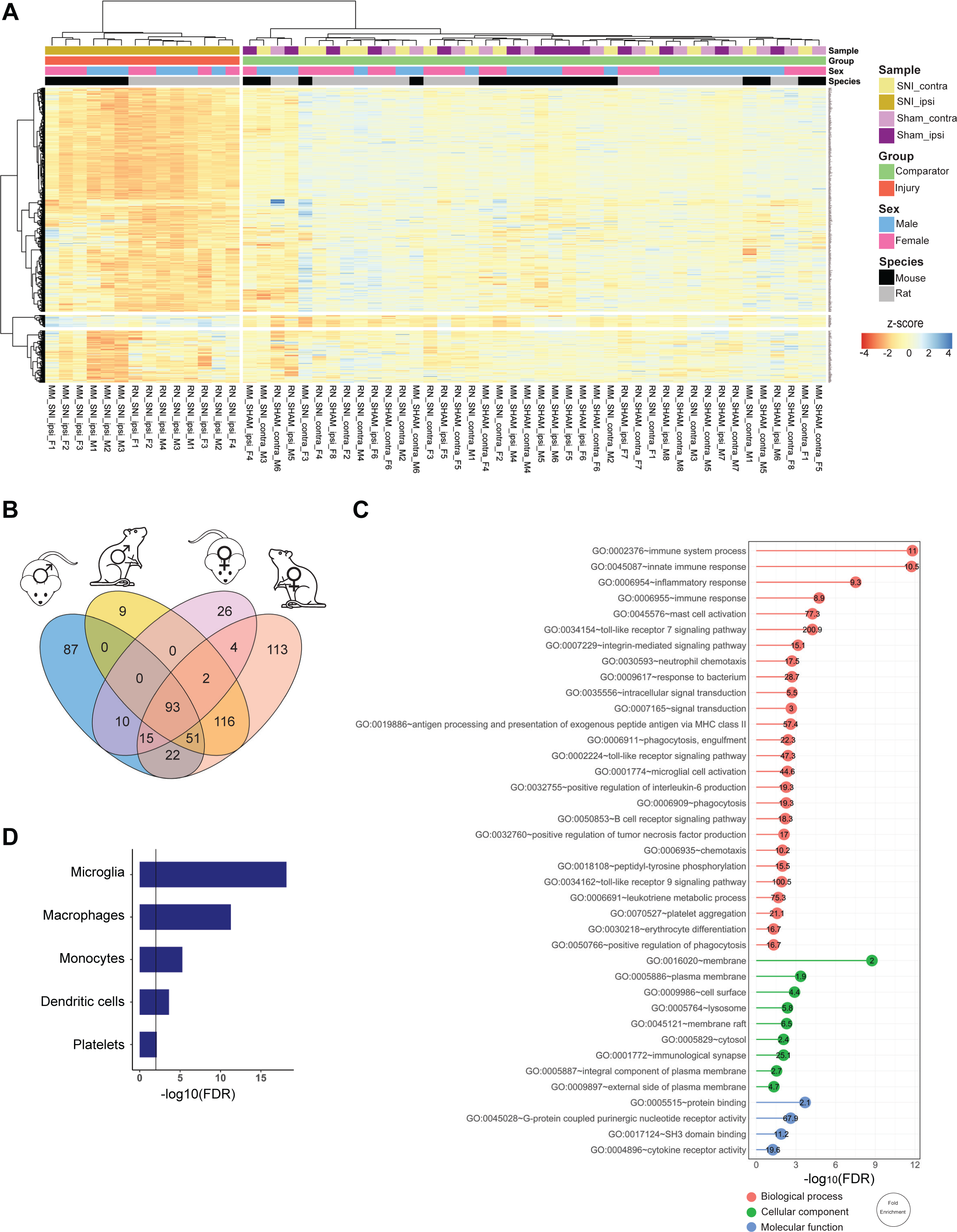
Peripheral nerve injury induces an immune response in rodents. (A) Heatmap showing the expression of the genes that were differentially expressed in at least one out of four (male mice, female mice, male rat and female rat) datasets, z-scores were calculated within species. (B) Venn diagram represents the number of differentially expressed genes between datasets. (C) Gene ontology enrichment of 93 conserved genes, biological processes are shown in pink, cellular components are shown in green and, molecular function are shown in blue. (D) Bar chart represents deconvolution profile of conserved genes obtained by scMappR package.

For genes that were differentially expressed after SNI in mice we compared the change in the level of expression in males versus females (Fig. 2E) and found that the change in expression level in females was highly positively correlated with that in males (R_pearson_=0.68, p<2.2*10^-16^ ). As in mice, the expression levels of SNI-evoked DEGs in male and female rats showed a significant positive correlation (R_pearson_=0.96, p<2.2*10^-16^, Fig. 2F). Moreover, by comparing mice and rat datasets we found that gene expression changes were significantly correlated between the two species (R_pearson_=0.83, p<2.2*10^-16^, Fig. 2G). Taking these findings together we conclude that transcriptional reprograming in response to peripheral nerve injury is significantly conserved in both sexes and in both species.

### Validation of combined analysis by sex and species

Focusing on the genes that were differentially expressed in males and females of both species we detected 93 DEGs increased in SNI_ipsi vs comparators (Fig. 3 A-B and table S1). In separate cohorts of animals, the validity of the RNA sequencing was tested in four of these DEGs: three of these had not been previously linked to neuropathic pain (Rasal3, Ikzf1, and Slco2b1) and for P2ry12 (Table 1) which had been linked (*12*). For each of genes the relative expression level measured by qPCR did not differ across sex or species (fig. S4A) and there was statistically significant correlation between the relative expression level measured by qPCR and that by RNAseq (fig. S4B).

**Table 1-.**
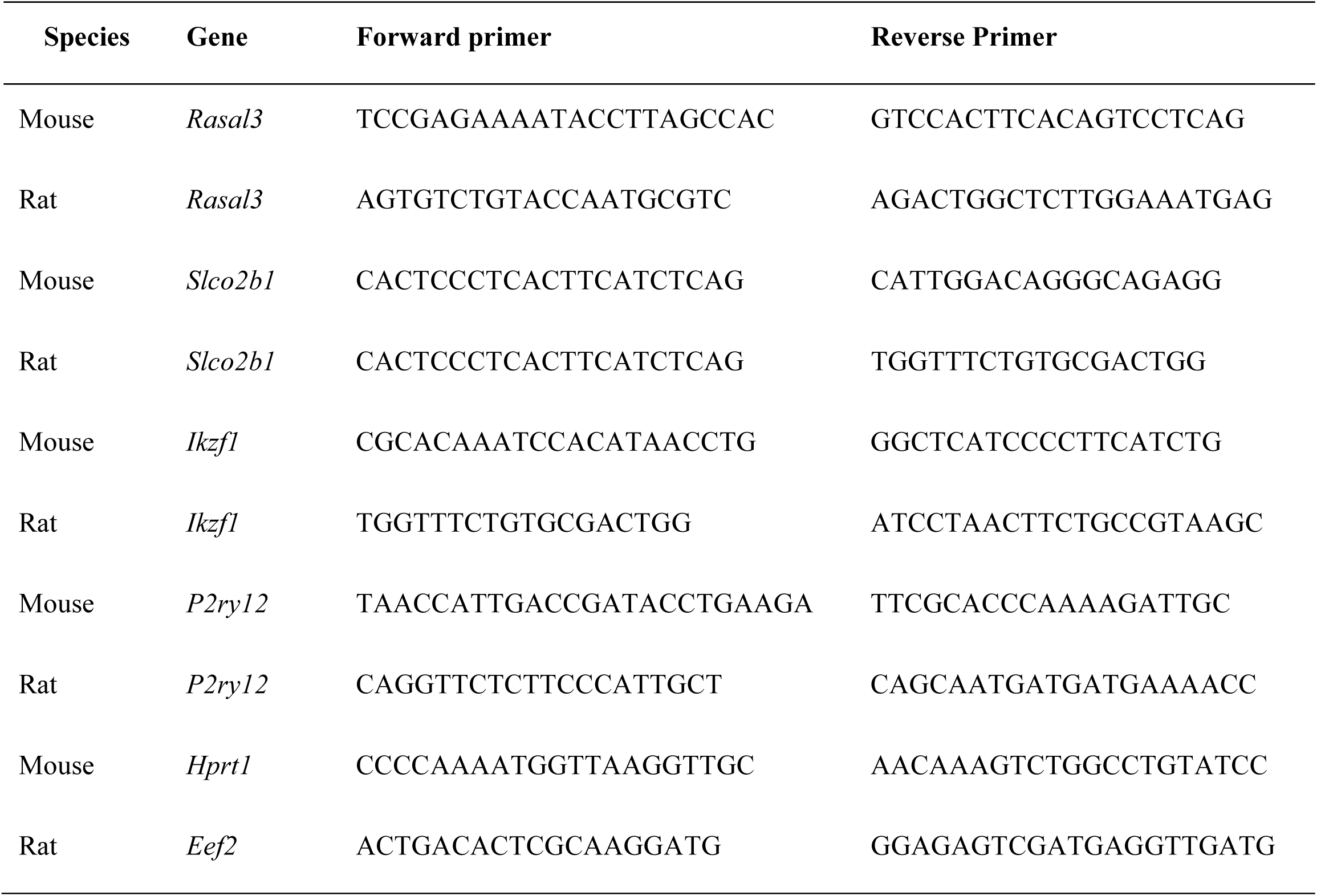
List of primers for candidate targets

The function of the 93 commonly upregulated genes was examined though Gene Ontology (GO) analysis (table S2). We found significant enrichment (false discovery rate (FDR) <0.05) of 26 biological processes of which 21 are directly related to immune responses (Fig. 3C). The predominant cellular components defined by GO analysis were related to membranes and those of GO molecular function were related to protein binding and G-protein coupled purinergic nucleotide receptor activity (Fig. 3C). To interrogate cell-type specificity for the common DEGs, we deconvolved the bulk RNA-seq dataset with scMappR (*13*) which uses publicly available single cell-RNAseq data from the Panglao database (*14*). Deconvolution analysis (Fig. 3D and table S3) revealed five cell types with FDR less than 0.05 with the top three being microglia cells (FDR=6.7*10^-19^, Odds Ratio= 20.1), macrophages (FDR= 5.1*10^-12^, Odds Ratio= 13.9), and monocytes (FDR= 15.8 *10^-5^, Odds Ratio= 8.56).

In exploring possible sex differences in the transcriptome changes induced by SNI we found 30 genes that were differentially expressed in female mice but not in males of either species and 117 genes that were differentially expressed in female rats but not in males (Fig. 3B). Of those female-specific DEGs four were common to females of both species including genes encoding neurofilaments light (Nefl), medium (Nefm) and heavy (Nefh) polypeptide and Proline-Serine-Threonine Phosphatase Interacting Protein 1 (Pstpip1). Notably, all of these genes were decreased following SNI. Gene ontology analysis and single-cell deconvolution for female mice and for female rats (fig. S5) revealed that while the individual transcripts differed there was a pattern common in both species that these DEGs were expressed in neurons. For males 87 genes were differentially expressed in male mice but not in females of either species one sex or species and we observed 9 genes that were differentially expressed in male rats but not in female rodents (Fig. 3B).

Together, the gene ontology analysis of the DEGs shows a pattern, biological processes, functions, cellular components and cell types, converging on microglia and immune response pathways in the dorsal horn ipsilateral to the nerve injury in both sexes and species, and at the same time the analysis reveals a female-specific pattern of DEGs conserved in both species. That there is a component of the transcriptional response of microglia genes which is conserved in both species and in both sexes, and that there is also a component of the response that shows sex differences in both species are consistent with transcriptional reprograming in the dorsal horn reported in the literature (*7, 8*) . Thus, we conclude that our approach of combining transcriptional profiles of sex and species together has face validity. By combining sex and species data we expected to have greater power than previous studies, and indeed we found changes in expression of genes for neuronal processes specifically in females, a finding not revealed by previous analyses.

### Defining a gene regulatory interactome network after peripheral nerve injury

We investigated whether there may be patterns in the repertoire of transcription factors regulating expression of the genes differentially expressed in the dorsal horn following nerve injury. To this end we used the ChEA3 database (*15*) which integrated six databases containing experimentally-defined transcription factor binding sites identified by chromatin-immune precipitation sequencing. We interrogated the ChEA3 database with the 93 conserved DEGs identified above. With the set of DEGs common across sex and species and with a cutoff of p<0.01 we identified 37 transcription factors (Fig. 4A, fig. S6 and Table.S4). Unsupervised hierarchical cluster analysis revealed 2 major clusters within the 93 conserved DEGs (Fig. 4B). Two of the transcription factors expressed in microglia that had not been previously linked to pain hypersensitivity are Lymphoblastic Leukemia Associated Hematopoiesis Regulator 1 (LYL1) and IKAROS Family Zinc Finger 1 (IKZF1). These transcription factors regulate 72% and 42%, respectively, of the common DEGs (Fig. 4C). We verified by PCR that expression of Ikzf1 was increased after SNI, in a new cohort of male and female mice and rat (fig. S4A).

**Fig 4.**
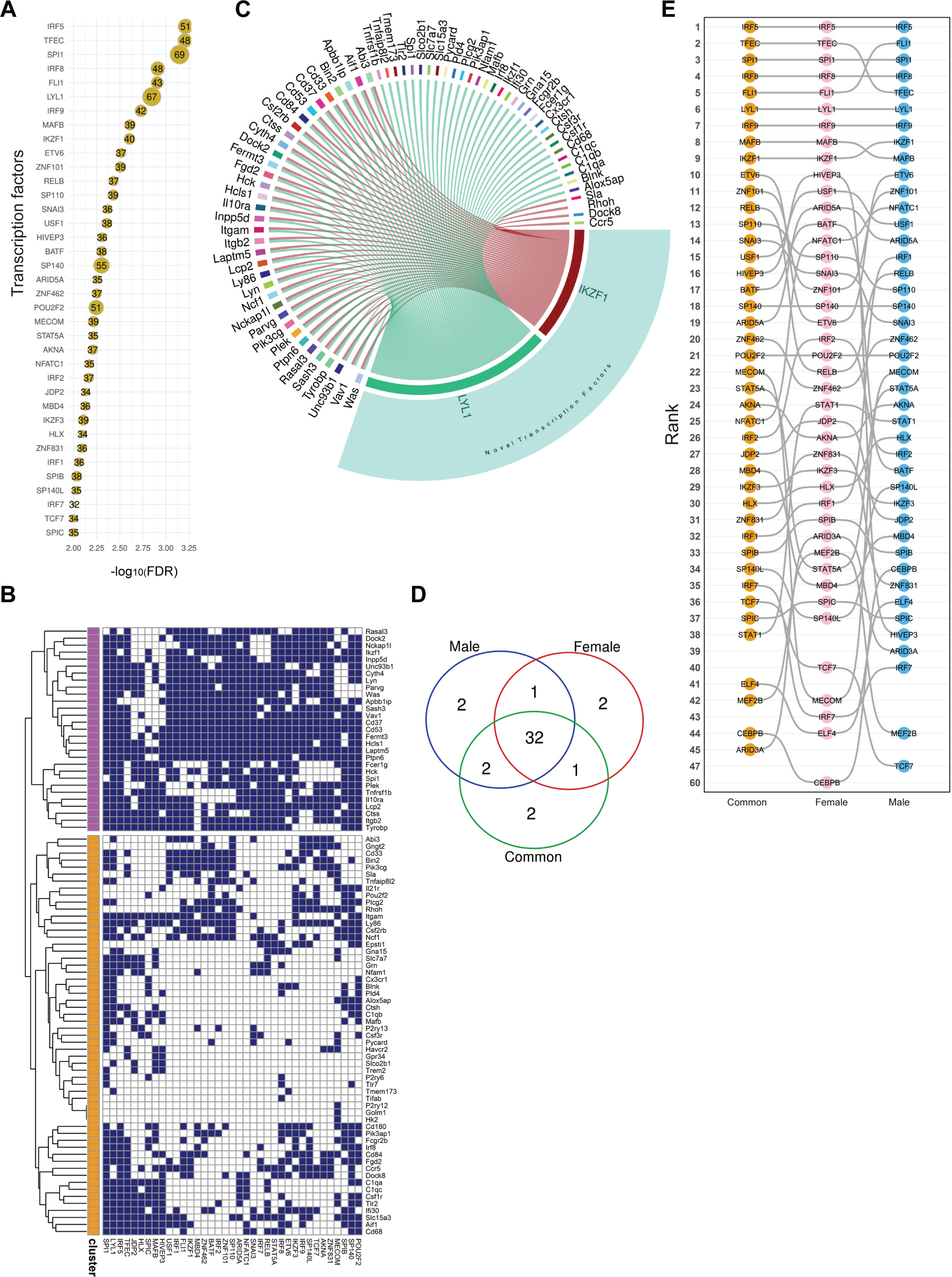
Gene regulation after peripheral nerve injury. (A) bubble plot showing transcription factors that can regulate conserved genes by ChEA3 database. (B) binary heatmap shows transcription factors and their targets. (C) Circoplot showing the relation between two novel transcription factors (LYL1 and IKZF1) and conserved genes. (D) Venn diagram represents the number of transcription factors between male, female and combined male and female datasets. (E) Bump chart visualizes the transcription factor ranking between three datasets of male, female and common (E).

We next analyzed transcription factor regulatory network in males and females separately. To investigate which transcription factors are contributing to the differential gene expression, we used the common DEGs in male of both species (n=144), and likewise in females, (n=114) (Fig. 4D). We found male and female rodents utilize transcription factors with different priority (Fig. 4E). For the lower ranked transcription factors there was increasing divergence in the rank order between males and females. We identified two male-specific (CEBPB, ELF4) and two female-specific (ARID3A, MEF2B) transcription factors. These transcription factors are reported to be expressed principally in microglia cells and in T cells, respectively (*16, 17*) (Fig. 4D, Table. S4).

### Targeting the sex- and species-conserved neuropathic pain interactome

As the DEGs and transcription factor networks in males and females were largely similar in both species, we wondered whether we could use the common DEGs to identify drugs that might reduce pain hypersensitivity in both sexes. From the proteins encoded by these DEGs we constructed a Protein-Protein Interaction (PPI) network using STRING (https://string-db.org) (*18*) This network was constructed with interaction scores greater than 0.9 and visualized in Cytoscape (*19*) (Fig. 5A, table S5). The resultant PPI network contained 38 nodes and 67 edges (interactions) which is significantly greater than predicted by a set of 93 proteins drawn randomly from the genome (PPI enrichment p-value < 1.0e−16). To identify the most influential nodes within the PPI network we calculated the Integrated Value of Influence (IVI) (*20*) for each node (table S6).

**Fig 5.**
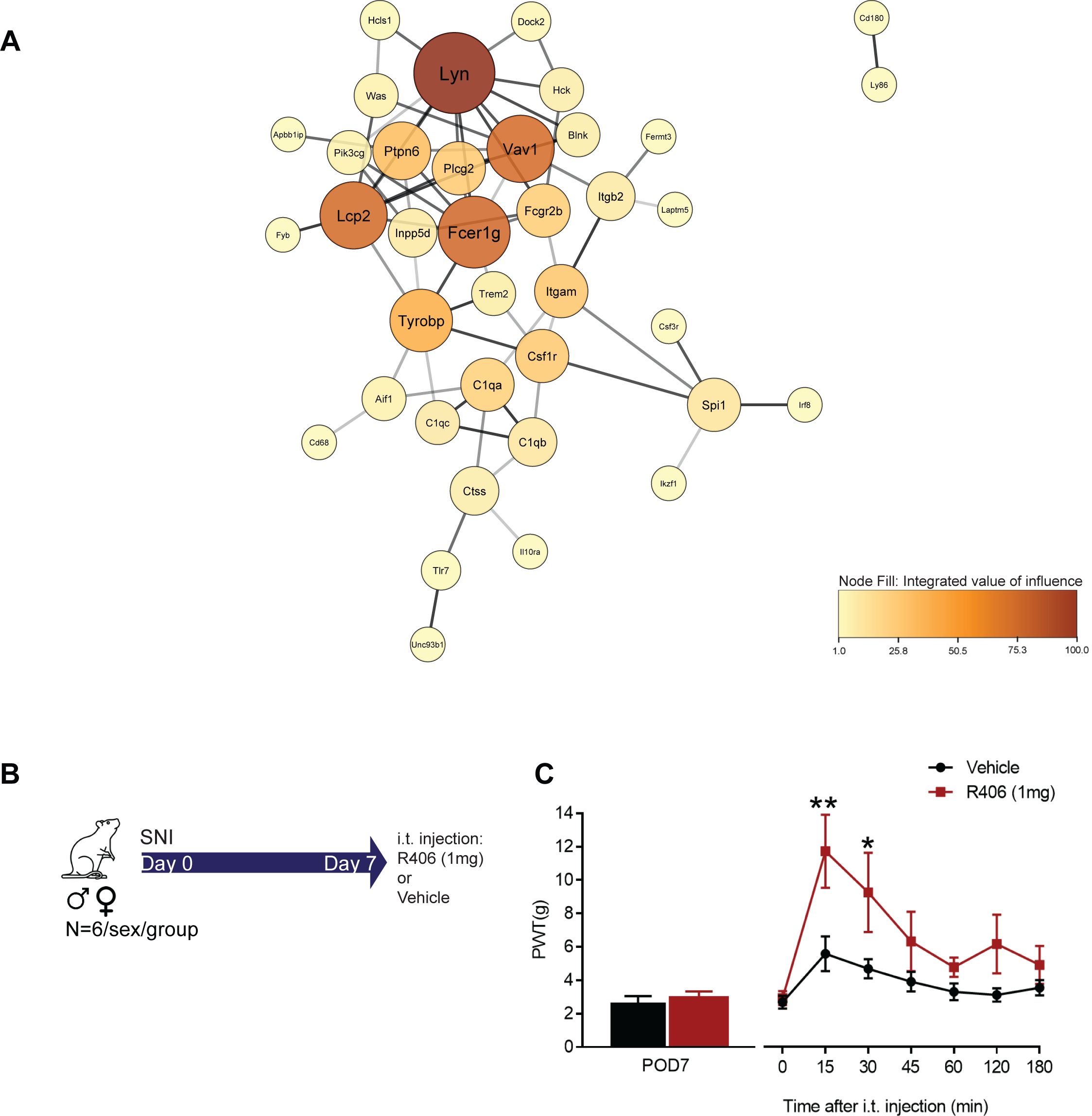
Targeting influential nodes inside conserved protein-protein interactome. (A) representing protein-protein interaction networks of conserved 93 DEGs retrieved from STRING database. This interaction map was generated using the maximum confidence (0.9). Color of the nodes is integrated the value of influence (IVI), Node size is relative to the node degree. Nodes without any connection are hidden from the network, edge thickness is based on evidence score. (B) Schematic diagram of experimental design for R406 *`in vivo`* trial. (C) Paw withdrawal threshold from von Frey filaments on the ipsilateral side 7 days after surgery in SNI animals, (N=6-7/sex/treatment) and comparing SNI ipsilateral of R406 (1mg) and vehicle. Comparisons were made by Bonferroni’s multiple comparisons test *p<0.05, **p<0.01. Data are mean ± SEM.

Separately, we interrogated the database of FDA-approved drugs – the Drug-Gene Interaction (DGIdb v4.1.0) (*21*) – with the list of 93 conserved DEGs. In the DGIdb we identified 186 drugs that affect one or more of the common genes (table S7). In order to find top FDA approved drugs that can target multiple influential nodes we calculated the Drug impact for each drug from the equation below (table S8).

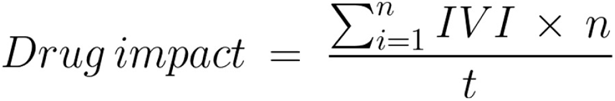

where *`n`* is representative of number of genes that are impacted by each drug and *`t`* is total number of nodes in the network. The five top-ranked were: 1-Fostamatinib, 2-Imatinib, 3-Bevacizumab, Daclizumab, Palivizumab, 4-Ibrutinib and 5-Etanercept. (Table 2).

**Table 2-.** list of top FDA approved drugs

| Rank | Top FDA Drug | Class | Targets in the network | Drug Impact |
| --- | --- | --- | --- | --- |
| 1 | Fostamatinib | Tyrosine kinase inhibitor | PIK3CG, HCK, LYN, CSF1R, CTSS, FCGR2B | 26.25 |
| 2 | Imatinib | Tyrosine kinase inhibitor | PIK3CG, HCK, LYN, CSF1R, IKZF1 | 18.14 |
| 3 | Bevacizumab | anti-vascular endothelial growth factor antibody | C1QA, C1QB, C1QC, FCGR2B | 6.63 |
|  | Palivizumab | Anti-respiratory syncytial virus F protein antibody | C1QA, C1QB, C1QC, FCGR2B |  |
|  | Daclizumab | CD25 antibody | C1QA, C1QB, C1QC, FCGR2B |  |
| 4 | Ibrutinib | Tyrosine kinase inhibitor | LYN, PLCG2 | 6.53 |
| 5 | Etanercept | Tumor necrosis factor alpha receptor inhibitor | TNFRSF1B, C1QA, FCGR2B, CD84, | 4.67 |

From this approach we predicted that drugs affecting the most influential nodes in the PPI network may inhibit pain hypersensitivity in both sexes. We tested this prediction for the top-ranked drug, fostamatinib. Fostamatinib is a pro-drug which yields the active molecule R406 by metabolism in the liver (*22*). We tested the effect of R406 in males and females seven days after SNI (Fig. 5C). Given that we implicated R406 from analyzing transcriptomes from the dorsal horn, we administered this drug intrathecally. We found that R406 significantly reversed SNI-induced mechanical hypersensitivity starting within 15 (p= 0.0016) and 30 mins (p= 0.0430) of the i.t. injection (Fig. 5D) with the effect in males indistinguishable from that in females (Fig. 5D and fig. S7). These findings are evidence confirming our prediction from the analysis of the PPI network and the DGIdb that a drug not previously associated with pain may reverse chronic pain hypersensitivity.

## DISCUSSION

Here, we generated a species-conserved, sex-conserved SNI-induced pain interactome network and, with an unsupervised approach, predicted safe therapies that might have the most impact in the interactome and thus might suppress pain hypersensitivity. We found that intrathecally administering R406, the active metabolite of the top-ranked FDA-approved drug fostamatinib, reversed mechanical hypersensitivity providing proof-of-concept to our approach. R406/ fostamatinib, which is clinically used to treat idiopathic thrombocytopenia purpura, was designed to suppress the kinase activity of spleen tyrosine kinase (Syk) (*23, 24*) making this kinase the most likely molecular target for the pain-reducing activity of this drug. We observed that Syk mRNA is substantially elevated in the ipsilateral dorsal horn by SNI providing a biologically plausible explanation for the effectiveness of R406. Moreover, the pain interactome includes upstream activators of Syk, Trem2 and CCR5, and downstream effectors in Syk signaling, VAV and PI3 kinase (Fig. 5D). R406 has been found to suppress the activity of a number of kinases and receptors (*25–28*) and thus a combined effect on multiple sites in the interactome network, in addition to its inhibition of Syk, may contribute to the analgesic action we discovered.

Syk is known to be expressed strongly in immune cells particularly macrophages, microglia, dendritic cells and B lymphocytes (*25*). The reversal of mechanical hypersensitivity by R406 in females as well as males may seem to suggest that the cell type affected by this drug is not microglia as interventions that suppress or ablate microglia differentially reverse pain hypersensitivity in males but not in females (*29*). This would be the case if R406 acts to suppress a pain-driving signal from microglia. But if R406 acts to induce microglia, or a subset thereof, to produce a pain-reducing signal then microglia could be the cell type in which R406 acts. Recently, a subtype of microglia, expressing cd11c, was reported to actively reverse hypersensitivity (*30*) in both sexes raising the possibility that R406 may act on this microglia subtype which strongly expresses Syk and for which the molecular signature gene, Itgax, is in the SNI-induced pain interactome (Fig. 5D). Alternatively, or in addition, meningeal macrophages, which are known to express Syk, have been implicated in controlling SNI-induced pain hypersensitivity (*16*) . While it appears that the most likely role for Syk, and hence the effect of R406, is in immune cells in the spinal cord, we cannot rule out an effect in neurons as a small proportion of three subtypes of excitatory neurons in the dorsal horn are reported to express Syk mRNA *`de novo`* after SNI (*16*) An effect of R406, directly or indirectly, on the cellular, neuronal processes of underlying SNI-induced pain hypersensitivity is consistent with the reported degeneracy of upstream immune cell signaling and the ultimate sex- and species-commonality of the principal pathological neuronal alterations, i.e. downregulation of the potassium-chloride cotransporter KCC2 and enhanced function GluN2B-containing NMDA receptors (*31*) .

From the 93 sex-conserved and species-conserved genes, the role of the proteins encoded by 17 of these genes in neuropathic pain has not been investigated to date (table S9). Based on gene ontology analysis, out of this 17 DEGs, Hck, Blnk, Sla, Lcp2 are involved in transmembrane receptor protein tyrosine kinase signaling pathway (table S10). the interaction of these genes and spleen tyrosine kinase needs to be further investigated.

In addition to defining the sex- and species-common genes, we explored the expression of genes for transcription factors that can regulate may regulate expression of these genes. We found that eight of the top 10 transcription factors have been linked to pain. Specifically, IRF5, the top-ranked transcription factor, is well-known to be markedly upregulated after peripheral nerve injury, and reducing expression of IRF5 prevents development of pain hypersensitivity in mice (*32, 33*). Two of the transcription factors we identified, Lyl1 and Ikzf1, have not been previously implicated in chronic pain hypersensitivity. Lyl1 is a basic helix-loop-helix (bHLH) type of transcription factor known to play a role on cell proliferation and differentiation and have a role on macrophages and microglia development (*34, 35*). IKZF1 is a type of lymphoid-restricted zinc finger transcription factor is known to regulate immune cells (*36*). It has been shown that Syk plays a crucial role for IKZF1 activation (*37*), therefore, R406 have a potential to disrupt IKZF1 nuclear localization and result in suppressing of IKZF1 targets.

The focus of the present paper on sex-conserved and species-conserved genes may seem contrary to a goal of considering sex as a biological variable in chronic pain (*38*). This focus was revealed by the results of our experimental and analytical design, and was only possible by examining both sexes, and both species, of rodents. It was only through testing and analyzing animals of both sexes that we were able to define those changes that are sex-different or sex-conserved without biasedly assuming that changes elucidated by studying only one sex, by far males, will generalize to the other sex. We did find sex differences in the transcriptional reprograming of the dorsal horn that were conserved in both rats and mice. Surprisingly, given past studies, we found evidence for differential cell type transcriptional changes induced by PNI linked to neurons. Specifically, the genes upregulated in female mice and rats were, to a first approximation, preferentially expressed in dorsal horn neurons. Exploring the role of the genes and gene networks discovered by this analysis therefore opens up the possibility of investigating the causal, i.e. necessary and sufficient, roles of proteins encoded by the genes we have identified as sex-specific. From our analysis it is apparent that transcriptional reprograming in the spinal dorsal horn in response to SNI has both sex-different and sex-conversed components.

In conclusion, we demonstrated that there is transcriptional reprograming in response to peripheral nerve injury that is conserved across sex and species. From deconvolving the species-conserved, sex-conserved pain interactome with the DGIdb database we created a ranking of FDA-approved drugs that we hypothesized may impact the pain interactome network. Given that the top hit, R406, pharmacologically inhibits Syk from humans and rodents (*23*), our discovery that this drug reverses SNI-induced mechanical hypersensitivity predicts that fostamatinib may reduce neuropathic pain humans, a prediction that is testable. We anticipate that our findings will provide a rational basis for speeding testing of potential analgesic agents, such as fostamatinib and others that impact the nerve injury-induced pain interactome, and therefore accelerate the pace of bringing new therapeutic options to those suffering with neuropathic pain.

## MATERIALS AND METHODS

### Study Design

Male and female C57BL/6J mice (n=6 per sex per condition aged 6-8 weeks) and Sprague Dawley rats (n=4 per sex per condition 7-8 weeks age) were purchased from The Jackson and Charles River laboratories at least two weeks before surgeries. All animals were housed in a temperature-controlled environment with ad libitum access to food and water and maintained on a 12:12-h light/dark cycle. In all experiments, animals were assigned to experimental groups using randomization. Experimenters were blinded to drugs and sex where possible; blinding to sex was ¬not possible in behavioural experiments. All experiments were performed with the approval of the Hospital for Sick Children’s Animal Care Committee and in compliance with the Canadian Council on Animal Care guidelines.

### Peripheral nerve injury

Neuropathic pain was induced in rodents using the spared nerve injury (SNI) model (Decosterd & Woolf, 2000). Briefly, animals were anesthetized with 2.5% isoflurane/oxygen under sterile conditions. An incision was made on the biceps femoris muscle’s left thigh and blunt dissection to expose the sciatic nerve. As a control, sham surgery was performed with all steps except sciatic nerve manipulation. The common peroneal and tibial nerves were tightly ligated and transected in the SNI model but left the sural nerve intact. The muscle and skin incisions were closed using 6-0 vicryl sutures in both groups.

### Tissue collection, library preparation and RNA sequencing

Animals were euthanized, and the L4-L5 lumbar dorsal horn of the spinal cord was harvested postoperative day 7 to study transcriptional changes. RNA was extracted from the tissue and preserved in RNALater (Invitrogen), and the library was prepared and sequenced using Illumina HiSeq 4000 by TCAG at The Hospital for Sick Children. The filtered reads are aligned to a reference genome using STAR (*39*). The genome used in this analysis was Mus musculus (GRCm38-mm10.0) and Rattus Norvegicus assembly (Rnor_6.0) after quality control, we calculated log2(CPM) (counts-per-million reads), and ran principal component analysis The differential gene expression analysis is done using DESeq2 (*40*) and edgeR (*41*) Bioconductor packages. Genes with adjusted p-Value <0.01 and fold changes greater than |0.5| were defined as differentially expressed genes (DEGs). In this study total of 24 samples from mice and 32 samples from rats were analyzed. We used three control groups (Sham_ipsi, Sham_contra and SNI_ipsi) as a reference to find differential expressed genes.

### Exploratory Analysis

Unsupervised hierarchal clustering was done by Euclidean method, number of optimal clusters were calculated using Elbow method in R. Enrichment analysis was performed on the DEG list using the Functional Annotation Tool in the DAVID website (https://david.ncifcrf.gov/) The protein-protein interaction (PPI) network of the proteins encoded by the DEGs was investigated using STRING v11.0 (*18*) to visualize protein-protein interaction. We used Cytoscape (*19*) Interactions with a score larger than 0.9 (highest confidence) were selected to construct PPI networks. Single edges not connected to the main network were removed. Transcription Factor enrichment analysis was performed using ChEA3, a comprehensive curated library of transcription factor targets that combines results from ENCODE and literature-based ChIP-seq experiments (*15*). Deconvolution of bulk RNA seq into immune cell types was evaluated using scMappR (*13*). The Drug Gene Interaction Database (DGIdb v4.1.0, www.dgidb.org) has been used to predict potential therapy for pain interactome (*21*) The integrated value of influence (IVI) was calculated by Influential R package (*20*). The impact of the drugs was calculated based on equation below:

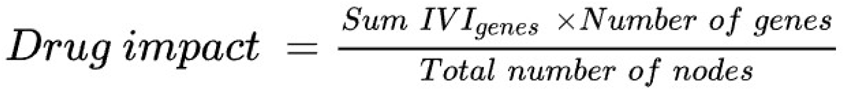

### Quantitative real-time reverse transcription-polymerase chain reaction

RNA was isolated by digesting L4:L5 spinal cord tissues in TRIZOL (Life Technologies) and cDNA synthesized using the SuperScript VILO cDNA kit (Life Technologies). qPCR was performed for 40 cycles (95 °C for 1 s, 60 °C for 20 s). Levels of the target genes were normalized against the average of four housekeeping genes (Hprt1 in mice and Eef2 in rats) and interpreted using the ΔΔCt method.

### Drug

R406 were purchased from Axon Medchem LLC (R406-1674). It was dissolved in DMSO, and corn oil Doses were determined in pilot experiments. Seven days post-SNI, rats were removed from their cubicles, lightly anesthetized using isoflurane/oxygen, and given intrathecal injections of R406 (1mg), in a volume of 20ul by 30-gauge needle.

### Behavioural test

Animals were randomized in experimental groups and behavioural experimenter was unaware of the treatment or design of the study. The mechanical withdrawal threshold of animals was tested on the ipsilateral paw using calibrated von Frey filaments of increasing logarithmic nominal force values. Animals were placed in custom-made Plexiglas cubicles on a perforated metal floor and were permitted to habituate for at least one hour before testing. Filaments were applied to the perpendicular plantar surface of the hind paw for one second. A positive response was recorded if there was a quick withdrawal, licking, or shaking of the paw by the animal. Each filament was tested five times with increasing force filaments (1-26g) used until a filament in which three out of five applications resulted in a paw withdrawal or when the maximal force filament was reached. This filament force is called the mechanical withdrawal threshold. The behavioural data is normalized as either percentage of baseline or presented as percent hypersensitivity.

### Statistical analysis

RNA-seq datasets were analyzed in R studio. For behavioral and Realtime PCR data, datasets were tested for normality using the Shapiro-Wilk test. qPCR data analyzed with the “pcr” R package, and behavioral data were analyzed by GraphPad Prism 9.3.1. One-way analysis of variance (ANOVA) or Kruskal-Wallis test was performed when comparisons were made across more than two groups. Two-way ANOVA (Bonferroni’s multiple) was used to test differences between two or more groups. T-test was performed to test differences between two groups. Statistical significance refers to *p< 0.05, ** p< 0.01, *** p< 0.001 Data are presented as mean ± SEM.

## Acknowledgments

Authors would like to thank Dr. David Finn and Dr. Katherine Halievski, Vivian Wang, Sofia Assi for technical assistance. This research was supported by a grant from CIHR (FDN-154336) to MWS. MWS held the Northbridge Chair in Paediatric Research. SG was supported by doctoral completion award and Massey College SAR, MMM was supported by a Pain Scientist Award from the University of Toronto Centre for the Study of Pain, and by a Restracomp postdoctoral fellowship from The Hospital for Sick Children Research Training Centre. Author contributions: MWS, SG conceived the project; MWS, MB supervised the research; SG and YT designed the experiments, MM, SG, YT designed in-vivo Fostamatinib experiment, SG, MM, YT, MK collected data and executed in vivo experiments, SG, AKR performed bioinformatic analysis, AS, AKR assisted with study design and interpretation of results. SG and MWS and wrote the manuscript with input from all authors. Competing interests: Authors declare that they have no competing interests. Data and materials availability: Data supporting the findings of this study are available within the article and its Supplementary material files and from the corresponding author upon reasonable request.

## Supplementary Materials

### Supplementary Figures

**fig. S1-.**
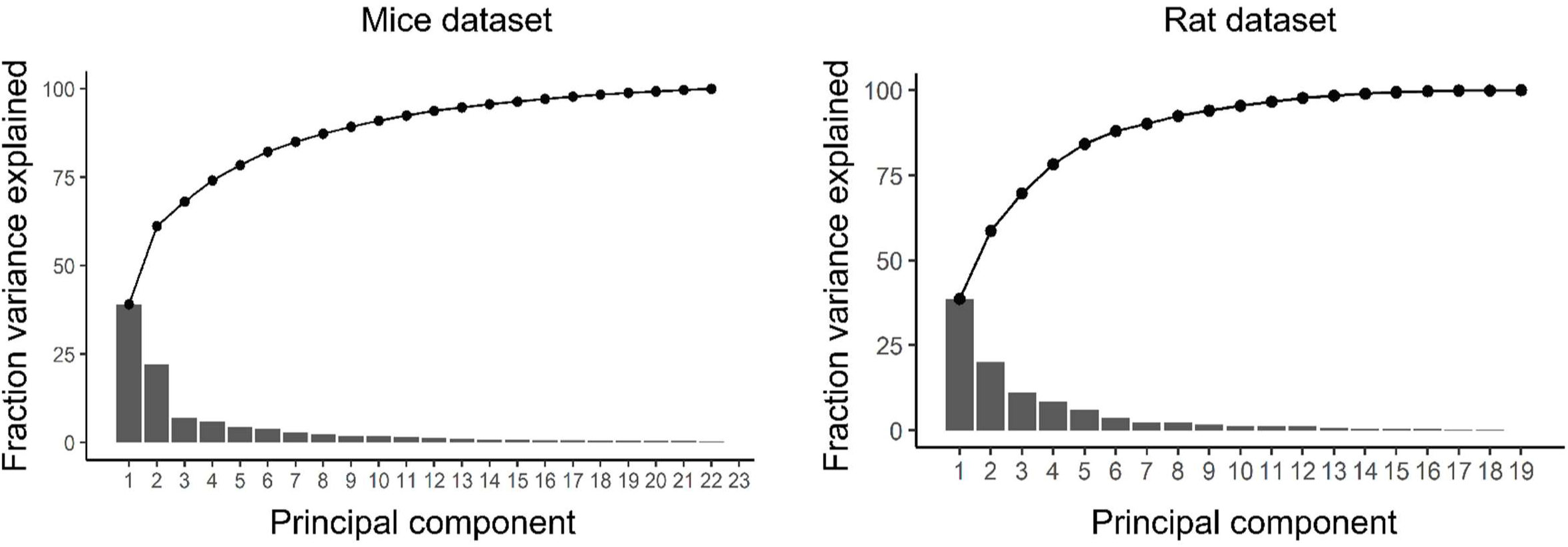
Principal components of mouse and rat datasets. Principal components, and explained variance from principal component analysis for mouse and rat datasets.

**fig. S2-.**
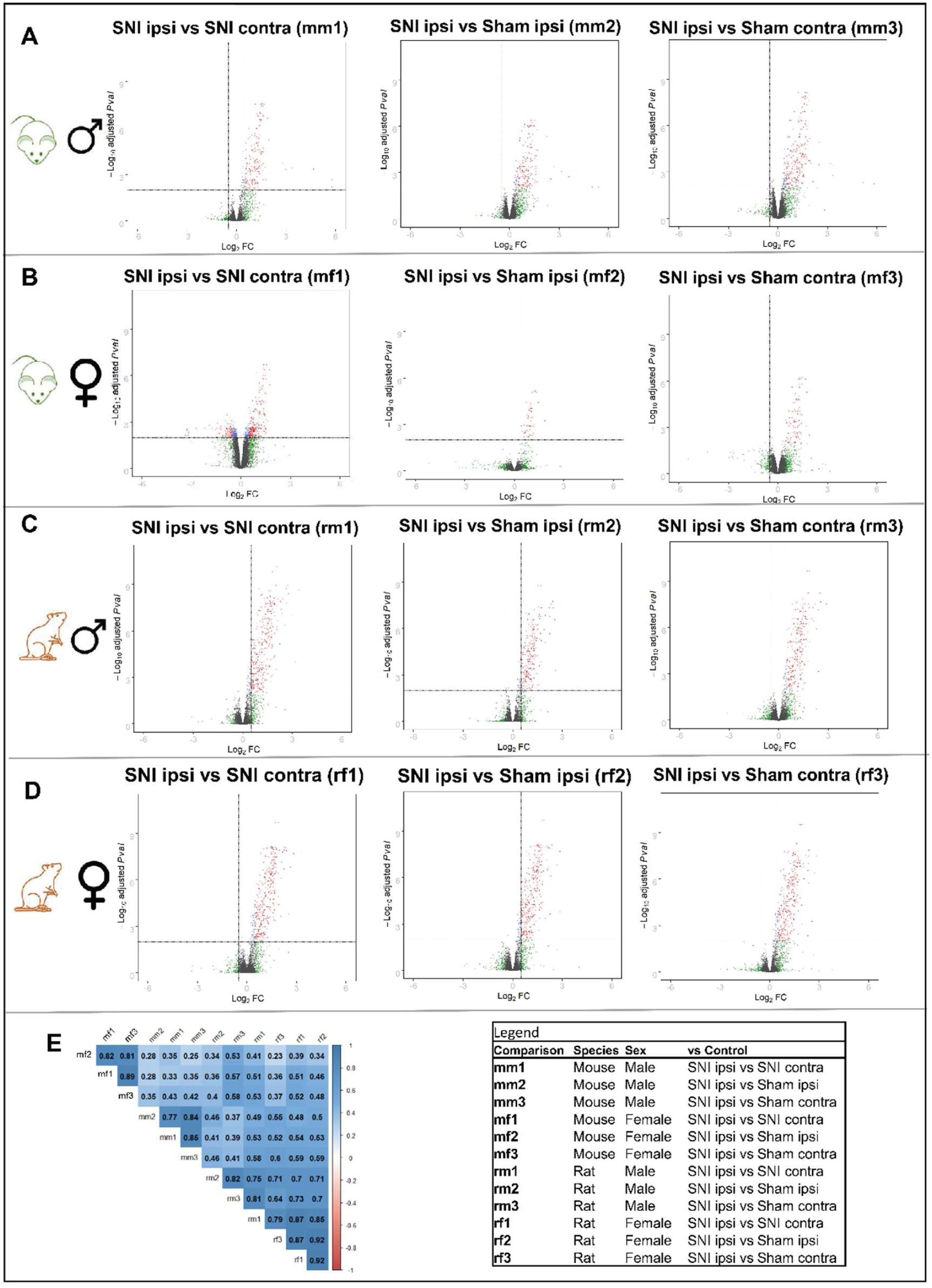
Pairwise comparison of SNI_ipsi vs each of the comparators. (A-D) Volcano plots of twelve pairwise comparisons (A) male mice (B) female mice (C) male rats and (D) female rats. (E) Correlation coefficients by Pearson method between 12 comparisons.

**fig. S3-.**
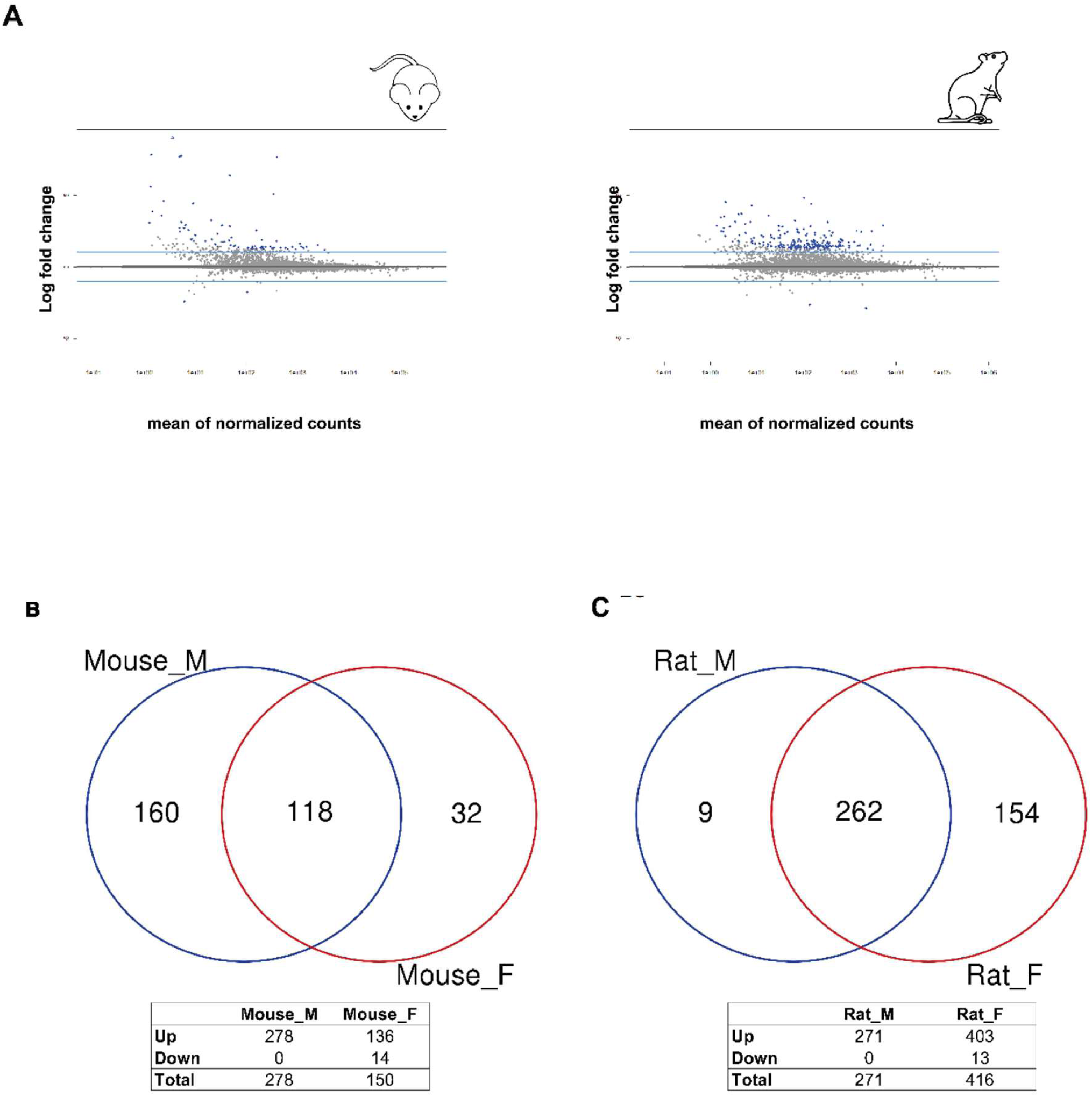
Overview of differential gene expression in mice and rat datasets. (A) MA plots of injured vs not injured sex combined. (B) Venn diagram shows common DEGs between males and females in mice and (C)Venn diagram for rat.

**fig. S4-.**
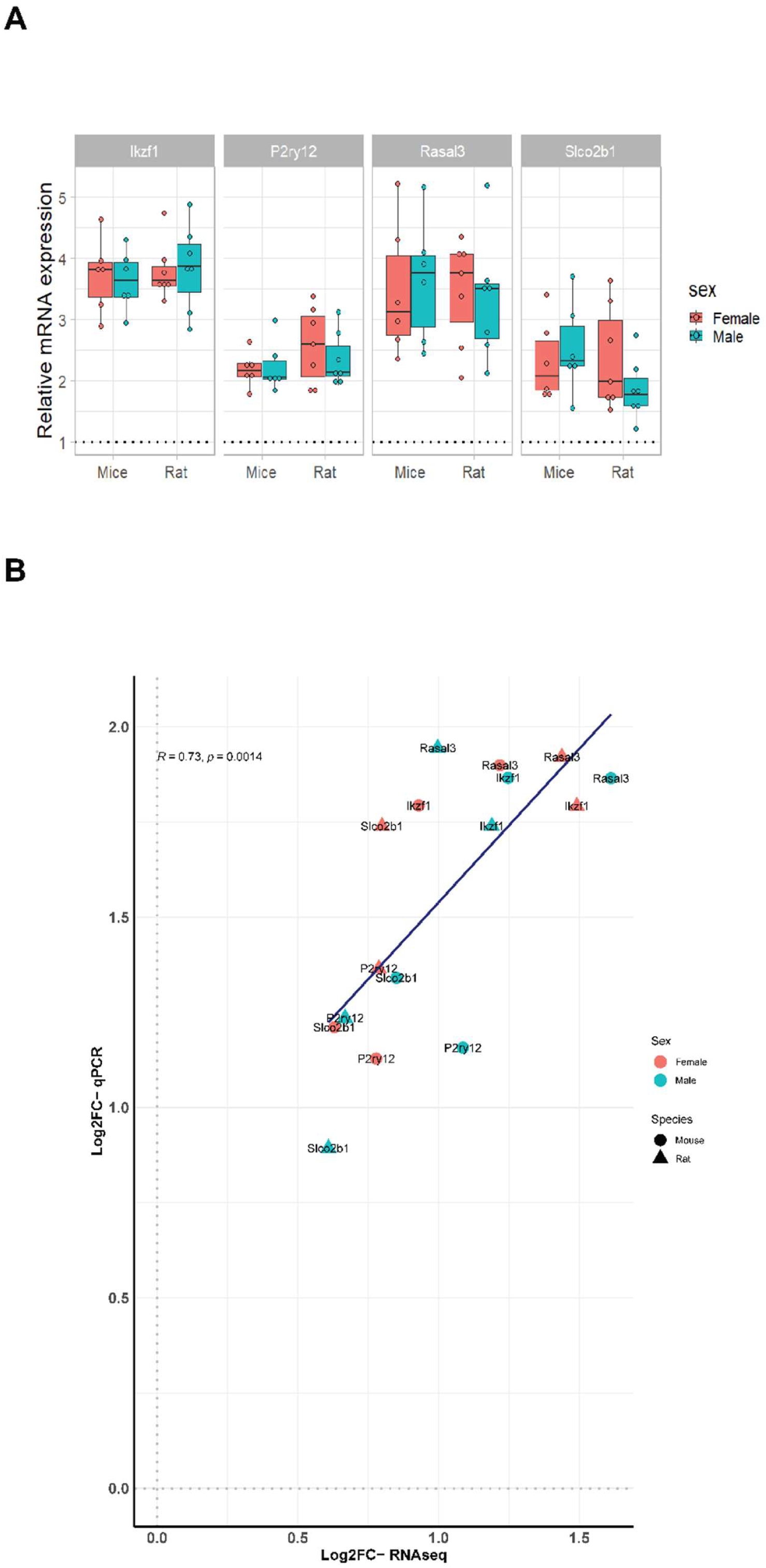
RNA-seq validation. (A) mRNA expression of 4 DEGs by qPCR, the delta-delta CT method was used to calculate the fold change vs SNI contra, the values represent the individual animal. (B) Pearson correlation of RNA-seq results and qPCR.

**fig. S5-.**
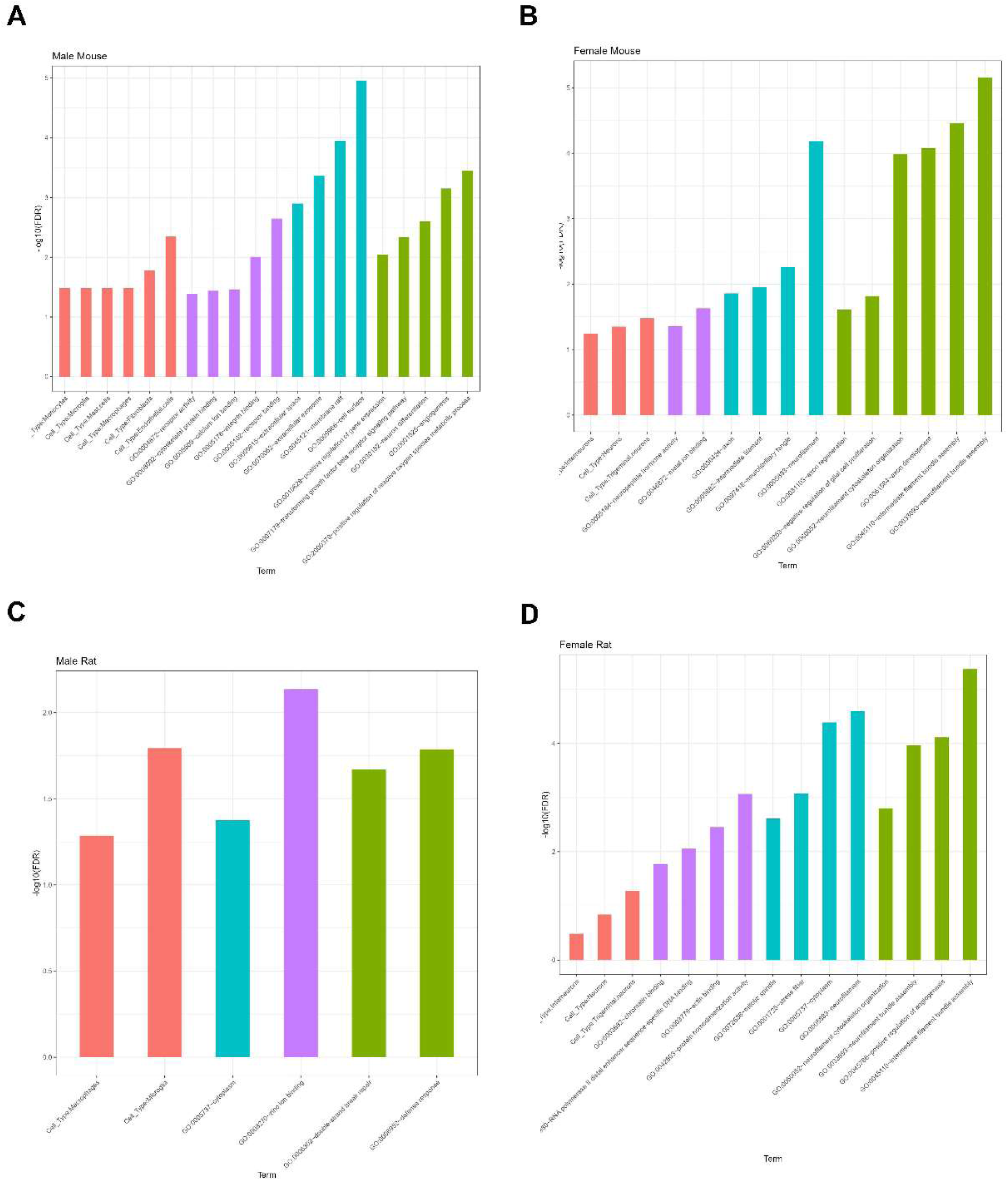
Gene ontology analysis and cell type profile for DEGs exclusive to sex or species. green color represents biological process, blue: cellular component, purple: molecular function is and red color represents cell-deconvolution. (A) male mice, (B) female mice, (C) male rats (D) and female rats.

**fig. S6-.**
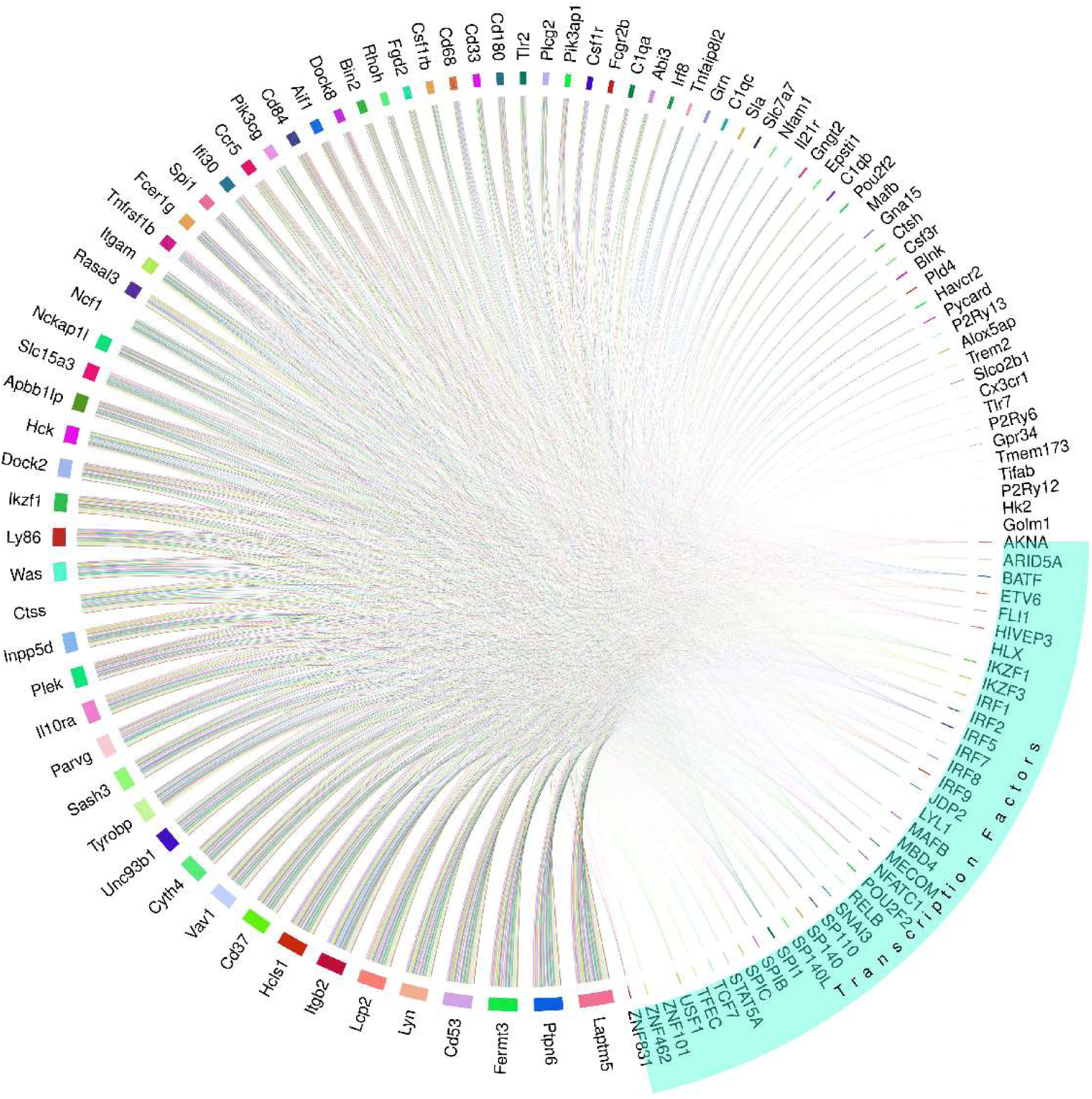
Gene regulation of conserved genes. Circoplot represents transcription factors and their connection with common differentially expressed genes.

**fig. S7-.**
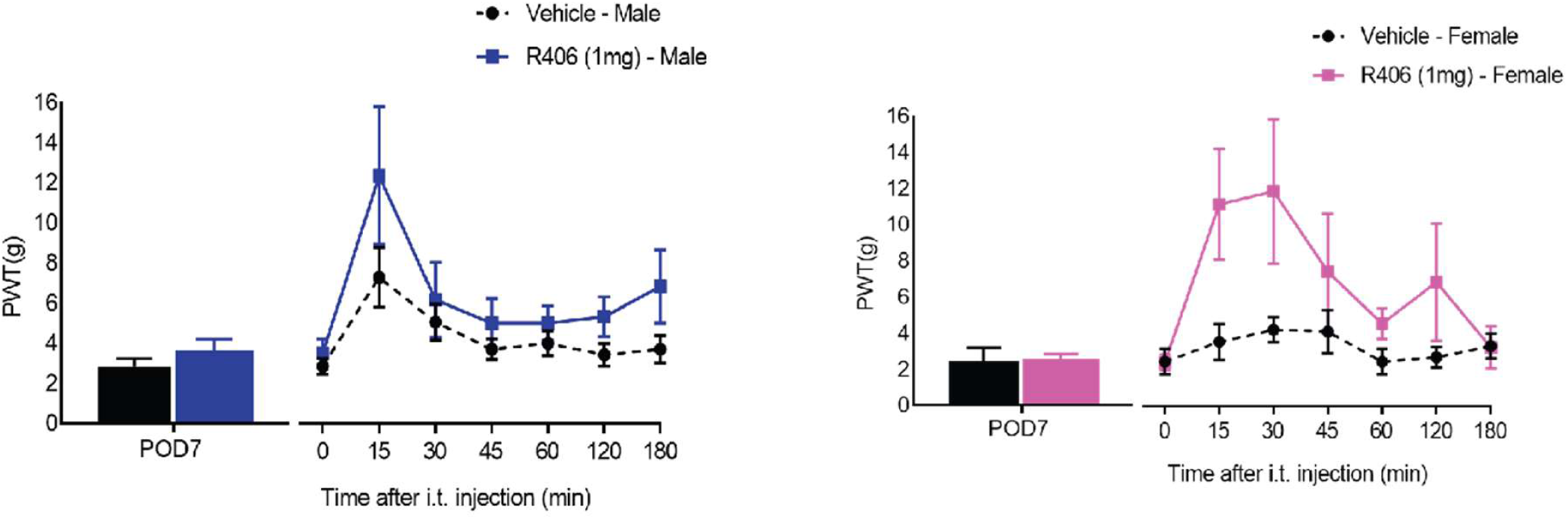
R406 efficacy in male and female rats. Paw withdrawal threshold from von Frey filaments on the ipsilateral side 7 days after surgery in SNI animals, (N=6-7/sex/treatment) and comparing SNI ipsilateral of R406 (1mg) and vehicle.

### Supplementary Tables

**table S1.** Differentially expressed genes between four datasets.

**table S2.** Gene Ontology results for conserved genes.

**table S3.** Single cell deconvolution of conserved genes

**table S4-** List of transcription factors regulating conserved genes within sex and species

**table S5-** Protein-Protein interaction network

**table S6-** Integrated Value of Influence for conserved nodes

**table S7.** Drug interaction with conserved genes

**table S8-** Drug impact calculation on network

**table S9-** List of novel targets

**table S10-** Gene ontology analysis for 17 novel genes

## Abbreviations

CNS: Central Nervous System
SNI: Spared Nerve Injury
Ipsi: Ipsilateral
Contra: Contralateral
DEG(s): Differentially Expressed Gene(s)
PC: Principal Component
i.t.: Intrathecal
IVI: Integrated Value of Influence
PPI: Protein-Protein-Interaction
DGIdb: Drug Gene Interaction Database
SYK: Spleen tyrosine Kinase

